# Vocal traits of shorebird chicks are related to body mass and sex

**DOI:** 10.1101/2021.12.30.474581

**Authors:** Kristal N. Kostoglou, Edward H. Miller, Michael A. Weston, David R. Wilson

## Abstract

Acoustic communication is critical during early life phases in precocial birds. For example, adult alarm calls can elicit antipredator behaviour in young, and chick vocalisations can communicate information to parents about chick identity, condition, location, sex, or age. We investigated whether chick calls of two species of Australian Charadriidae vary with sex or body mass. We handled Red-capped Plover *Charadrius ruficapillus* and Southern Masked Lapwing *Vanellus miles novaehollandiae* chicks for purposes of measurement, blood sampling, and banding. We opportunistically recorded their distress calls while in the hand, and analysed the calls to determine whether call structure is related to sex or body mass (a proxy for age). We measured five traits per call, plus time intervals between successive calls, for 26 plover chicks (2600 calls) and 95 lapwing chicks (6835 calls). In plovers, inter-call intervals were shorter in males and both inter-call interval and the dominant frequency range of calls decreased with increasing body mass. In lapwings, frequency modulation (computed as the range in the rate of change of the dominant frequency) was lower in male calls. The dominant frequency range of lapwing calls decreased with mass in both sexes, but the decline was greater in males, resulting in a lower dominant frequency range in males. Frequency modulation and entropy of lapwing calls also decreased with increasing body mass. Minimum dominant frequency did not change with body mass or sex in either species. Our study provides the first evidence for charadriid chicks of (a) a sexual difference in call structure and rate and (b) gradual growth-related changes in call structure and rate, across chicks. Studies on calls from a greater range of chick ages and from more species within this large and diverse family would be valuable. We provide a foundation for further studies of shorebird vocalisations during growth, which may elucidate the development and functional significance of such vocalisations.

---

Acoustic communication can be critical in the early life of birds. In nestlings, begging calls can solicit parental feeding, contact calls can help parents locate and identify offspring, and distress calls given during social separation, environmental challenges, or handling can startle predators or elicit help from parents, conspecifics, or other nearby species (Stefanski and Falls 1972, Sethi et al. 2012). Several studies have investigated the function of calls in juvenile birds (e.g. Desmedt et al. 2020) but few have considered the structure or development of juvenile calls. Developmental changes in chick vocalisations can allow parents to assess chick condition (e.g. Goedert et al. 2014) and have been documented for some precocial species (e.g. Würdinger 1970, Desmedt et al. 2020). With some exceptions (Herting and Belthoff 2001, Odom and Mennill 2010), there is a general trend for the dominant frequency of calls to be inversely related to body mass or size across species, possibly due to the greater mass of vocal structures or respiratory muscles (Suther and Zollinger 2004). Larger birds also may be constrained in producing higher frequencies (e.g., via stronger constraints on tissue elasticity with larger anatomical structures; Demery et al. 2021). Intraspecifically, changes in vocalisations can occur gradually as body mass increases with age (Dragonetti et al. 2013), or in a step-like fashion (Klenova et al. 2014).

Vocal traits also can differ between sexes, and play important roles in reproduction, territorial defence, and other social activities (Buck et al. 2021). Sexual differences emerge at various times during development in different species, from several days old to after fledging (Cosens 1981, Saino et al. 2008) to as late as at sexual maturity (Klenova et al. 2014), but no general patterns are evident (Tikhonov 1986, ten Thoren and Bergmann 1987a,b, Volodin et al. 2015). The nature of sexual differences in calls also varies across species, for example, in the dominant frequency (ten Thoren and Bergmann 1987a), duration (Tikhonov 1986), or amplitude (Saino et al. 2008) of calls. Some interspecific variation in the occurrence and nature of sex-related differences may result from patterns of growth, sexual differences in body size, or ecological or social factors (Saino et al. 2003, Volodin et al. 2015, Austin et al. 2021).

Vocal communication in shorebirds can help keep precocial young close to their parents and away from predators, and might also play a role in mediating sexual differences in the survival, parental care, and dispersal of young (Pakanen et al. 2015, Eberhart-Phillips et al. 2017, Lees et al. 2018, 2019). However, despite a plethora of studies and reviews on social organisation, breeding biology, behavioural ecology, and communication in shorebirds, vocal development has been examined in only two species. Adret (2012) quantified age-related changes in measures of call frequency and noted a sexual difference in call rates for Pied Avocet *Recurvirostra avosetta* chicks. Dragonetti et al. (2013) described qualitative changes of Eurasian Stone-curlew *Burhinus oedicnemus* chick calls with age. To extend knowledge of vocal communication and development in shorebirds, we investigated potential relationships of call traits to body mass and sex in chicks of two shorebird species. When in the hand for banding and blood sampling, chicks of Red-capped Plover *Charadrius ruficapillus* and Southern Masked Lapwing *Vanellus miles novaehollandiae* (“plovers” and “lapwings” hereafter) often utter distress calls. We recorded these calls to document their structure (Miller et al. 2021), and to determine whether they change with body mass (a proxy for age) or differ between sexes. We predicted that call structure, especially frequency-related characteristics, would change with increasing body mass (Demery et al. 2021). In some precocial bird species, it would likely be beneficial for parents to distinguish the sex of their offspring, for purposes of parental defence and care (Barrios-Miller and Siefferman 2013, Lees et al. 2018). We therefore predicted sexual differences in chick calls. Our study provides the first account of call development for any species of Charadriidae.

## Methods

We studied plovers from October 2017 to March 2018 (Cheetham Wetlands, Victoria, Australia; 37° 54’ S 144° 47’ E; 420 ha). Male chicks have slightly longer tarsi than females (Lees et al. 2019), but otherwise the sexes are indistinguishable until their second immature plumage (Marchant and Higgins 1993). We studied lapwings, which have sexually indistinguishable young (Lees et al. 2018), from June to September 2018 (Phillip Island, Victoria, Australia; 38° 29’ S, 145° 14’ E; 10,000 ha). Parents brood and defend their chicks which attain the capacity for sustained flight at approximately 35 (plovers) and 45 (lapwings) days of age (Temple-Smith 1969, Marchant and Higgins 1993, Lees et al. 2018).

We conducted extensive searches for nests from vehicles and by foot. Upon discovering a nest, we estimated approximate hatching dates by floating the eggs (Liebezeit et al. 2007), assuming incubation periods of 30 days (plovers) and 32 days (lapwings) (Marchant and Higgins 1993, Lees et al. 2018). We returned to nests around our estimated hatching dates to capture chicks. Occasionally, we captured older unbanded chicks away from known nests. For each captured chick, we measured body mass (Pesola spring balance: ± 0.1 g), and tarsus, tarsus plus toe, culmen, and head plus culmen lengths (dial calliper: ± 0.1 mm) (Rogers et al. 1990). All body measurements are provided within Supporting Information Tables S1 and S2. We use only body mass as a proxy for age because: 1) we did not know exact ages of most chicks, 2) all variables describing body size were highly inter-correlated (*r*_Pearson_ ≥ 0.75 for all pairwise combinations), and 3) a relationship of call structure to body mass occurs in other species (Martin et al. 2011). We also obtained ca. 50 μL of blood from the tarsal vein of each chick, which permitted genetic sexing (DNA Solutions™). We recaptured and measured (excluding blood extraction) five plover chicks (mean interval between captures: 8.6 ± 3.4 days, range: 6–14 days, *n =* 5) but did not recapture any lapwing chicks. We cite mean ± SD throughout.

### Call recording and acoustic analysis

We processed chicks singly (plover: 9.8 ± 5.1 minutes, *n* = 26; lapwings: 5.7 ± 2.9, *n* = 95) in a quiet, sheltered location. We recorded their vocalisations using a portable digital recorder (Roland R-26, WAVE format, 44.1 kHz sampling rate, and 16-bit depth) and an omni-directional Sennheiser ME 2-II microphone (50−18,000 Hz frequency response) held approximately 5 cm from the chick. Recordings sometimes included calls of siblings held nearby, but these calls were distinguishable in our recordings; we assume they do not influence the variables we measure. For the five plover chicks that were recaptured, we analysed vocalisations from their second capture.

We provide a detailed account of acoustic analysis in Supplemental Methods. Briefly, after filtering calls, we identified start and end times of each call and we divided each call into a series of contiguous 2.9-ms time bins. We measured the Shannon spectral entropy and dominant frequency from a mean power spectrum for each bin. We used six call traits for analyses: (1) call duration, the time interval between the start and end of a call (seconds); (2) inter-call interval (ICI), the interval between the end of a call and the start of the next (seconds); (3) entropy, the average of all spectral entropy values within a call (unitless); (4) minimum dominant frequency (kHz); (5) dominant frequency range, the difference between a call’s minimum and maximum dominant frequency (kHz); and (6) frequency modulation. For lapwings, we calculated frequency modulation by fitting a series of polynomial regressions (up to twelfth order) to the dominant frequency and time values of the call, selecting the best-fitting model (Supporting Information Fig. S1). For each 2.9-ms time bin, we calculated the slope of the tangent to the polynomial curve and used the range of slopes across the call as our measure of frequency modulation (kHz/s) (Supporting Information Fig. S1). For plovers, polynomial regression curves could not adequately model the frequency modulations. Instead, we defined frequency modulation as the cumulative absolute change in dominant frequency across all 2.9-ms time bins, divided by call duration (kHz/s). We also calculated maximum dominant frequency as a separate variable, but excluded it because it was highly correlated with dominant frequency range for both species (*r*_Pearson_ = 0.86 and 0.89 for plovers and lapwings, respectively) and because statistical models exhibited poor fit (see below). Correlations among the remaining six variables were all < 0.7. Not all variables could be calculated from every call.

### Statistical analysis

Separate generalised linear mixed models (GLMMs) examined possible relationships among call traits, body mass and sex for each species. We included chick identity nested within brood identity as a random effect to account for possible dependencies among calls recorded from the same chick or brood. We specified a Toeplitz covariance structure (Glaz and Yeater 2020) to account for sequential autocorrelation between calls. We included body mass and sex as main effects in all models. For some traits, an interactive effect between body mass and sex might occur, although we had no *a priori* reason to expect this. Nevertheless, for each analysis, we compared a model with main effects only to a model with main effects plus an interaction term between body mass and sex. We based model selection on the Akaike Information Criterion (AIC) values, with best models being identified where they differed from the alternative candidate model by ΔAIC > 2 (Symonds and Moussalli 2011). Where candidate models appeared to be equally informative (ΔAIC ≤ 2), we did not include the interaction term in the final model (the main-effects-only model always had the lowest AIC).

We tested model assumptions and evaluated model fit by comparing observed data to simulated data derived from the model (Hartig 2018). For lapwings, many calls had a measured frequency modulation of zero, which resulted in poor model fit (i.e. outliers had significant leverage) that could not be improved through the use of zero-inflated models (Brooks et al. 2017). We retained the zeros within the final model because they were not errors, and their removal did not change the results with respect to statistical significance (Supporting Information Fig. S2 and Table S3). When analysing entropy, we excluded calls for which the recording was saturated (the input of a signal was greater than the output, creating clipping and distortion to the shape of the waveforms; plovers, *n =* 939 calls; lapwings, *n =* 3647). Saturation can affect entropy, and the entropy differed significantly between saturated and non-saturated calls (Supporting Information Table S4). When analysing ICIs, we excluded recordings with < 10 calls (plovers, *n =* 3 recordings; lapwings, *n =* 20) because calling was too sporadic to get a reliable measure of ICI. We used the median ICI of each individual’s recording (excluding any sibling calls), since we had no objective way of distinguishing calling bouts.

Transformations of data were performed to improve distribution and model fit. We applied a reciprocal transformation on body mass for both species for all models. For lapwings, we added a constant (i.e. 2) and then logarithmically transformed the data for frequency modulation, dominant frequency range, and ICI. We performed the same transformation for dominant frequency range and ICI for plovers; however, as entropy was left-skewed, we subtracted values from a constant (2) and then log-transformed the data. We indicate probability distributions and link functions within the tables.

We performed analyses in R using the glmmTMB package for running mixed models, DHARMa for model assumption testing, bbmle for calculating ΔAIC values, psych for creating histograms and scatterplots, base R for creating and observing simulated data and ggplot2 and ggeffects for figures (Wickham 2016, Brooks et al. 2017, Hartig 2018, Lüdecke 2018, Bolker and R Core Team 2020, 2021, Revelle 2021). We controlled the experiment-wise type I error rate by applying a Bonferroni correction, which adjusted our alpha value from 0.05 to 0.0083. For figures, ICIs, entropy, and dominant frequency range were back-transformed and presented on the original scale in figures. Estimated marginal means (hatched and solid lines) and 95% confidence intervals (grey shading) were calculated using ggeffects (Lüdecke 2018).

## Results

For plovers, we recorded 2600 vocalisations from 26 individual chicks (1–248 calls per chick; 570 calls from 9 females and 2030 calls from 17 males), over the body mass range of 3.5–20.5 g (8.5 ± 4.6 g). This range corresponds to chick ages from the day of hatching to approximately 4 weeks of age (Lees et al. 2019).

For lapwings, we recorded 6835 vocalisations from 95 individual chicks (1–336 calls per chick; 3174 calls from 46 females and 3661 calls from 49 males), over the body mass range of 15.2–177.0 g (43.6 ± 30.2 g). This range corresponds to chick ages from the day of hatching to approximately 5 weeks of age (Thomas 1969, Moffat 1981).

For plovers, ICIs were shorter for males than for females and, as mass increased, ICIs and the dominant frequency range of calls decreased (Table 1, Fig. 1). There were no significant relationships of call duration, entropy, minimum dominant frequency, or frequency modulation to sex or body mass.

**Table 1.**
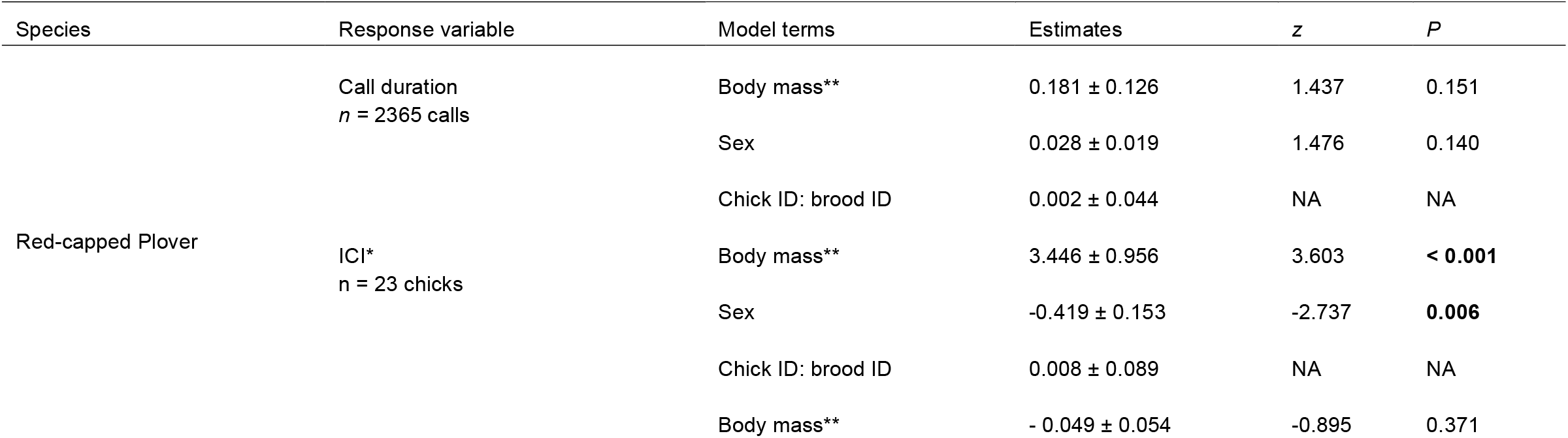
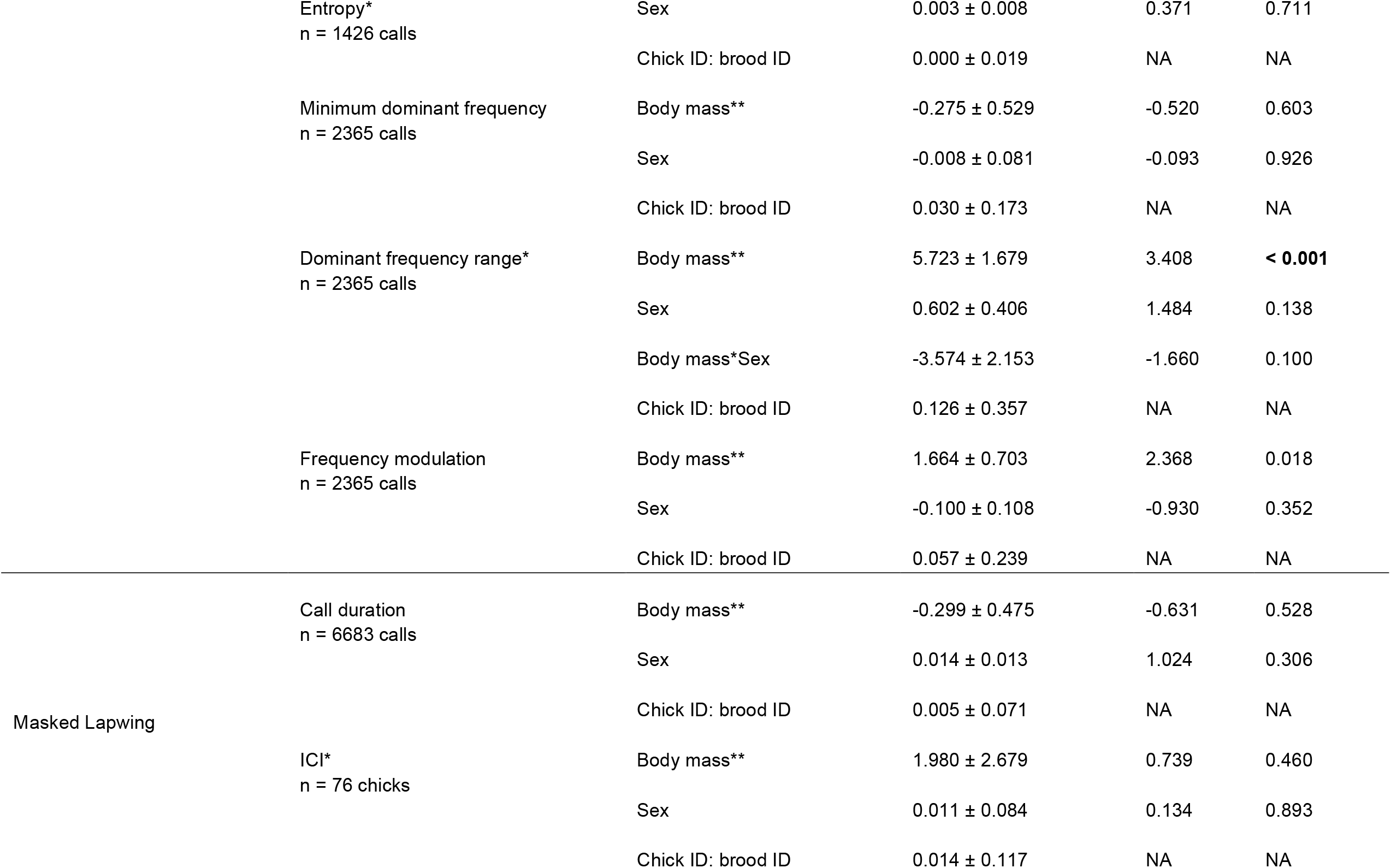
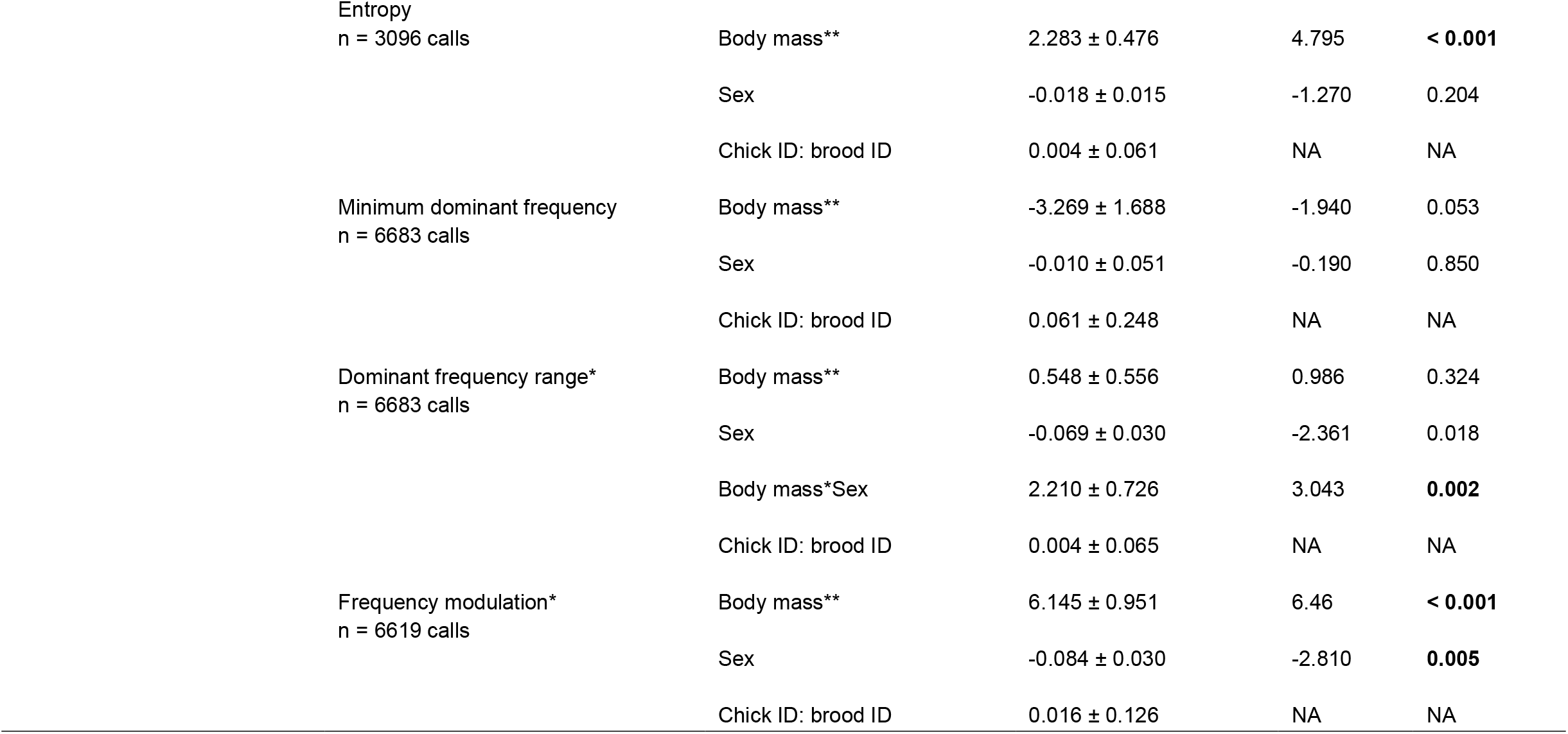
Results of separate generalised linear mixed models investigating possible relationships between body mass (reciprocal transformation) and sex (the interaction was selected for inclusion for the frequency range model only) with call duration, inter-call interval (ICI), entropy, minimum frequency, frequency range, and frequency modulation for plovers and lapwings. Chick identity nested within brood identity was included as a random effect for all analyses. We specified a Toeplitz covariance structure for all models. The reference category for sex was female. Estimates are presented as estimates of coefficients ± standard error for fixed effects, and variance ± standard deviation for the random effect of identification nested within brood identity. The probability distribution was Gaussian and we used an identity link function for all variables except frequency modulation (plovers), for which we used a negative binomial (linear parameterisation) distribution with a log link. * = logarithmically transformed data. ** = to aid interpretation, note that the reciprocal transformation reflects the sign of coefficients.

| Species | Response variable | Model terms | Estimates | *z* | *P* |
| --- | --- | --- | --- | --- | --- |
| Red-capped Plover | Call duration<br>*n* = 2365 calls | Body mass** | 0.181 ± 0.126 | 1.437 | 0.151 |
| | | Sex | 0.028 ± 0.019 | 1.476 | 0.140 |
| | | Chick ID: brood ID | 0.002 ± 0.044 | NA | NA |
| | ICI*<br>*n* = 23 chicks | Body mass** | 3.446 ± 0.956 | 3.603 | **< 0.001** |
| | | Sex | -0.419 ± 0.153 | -2.737 | **0.006** |
| | | Chick ID: brood ID | 0.008 ± 0.089 | NA | NA |
| | | Body mass** | - 0.049 ± 0.054 | -0.895 | 0.371 |

| Species | Response variable | Model terms | Estimates | z | P |
| --- | --- | --- | --- | --- | --- |
|  | Entropy*<br>n = 1426 calls | Sex | 0.003 ± 0.008 | 0.371 | 0.711 |
|  |  | Chick ID: brood ID | 0.000 ± 0.019 | NA | NA |
|  | Minimum dominant frequency<br>n = 2365 calls | Body mass** | -0.275 ± 0.529 | -0.520 | 0.603 |
|  |  | Sex | -0.008 ± 0.081 | -0.093 | 0.926 |
|  |  | Chick ID: brood ID | 0.030 ± 0.173 | NA | NA |
|  | Dominant frequency range*<br>n = 2365 calls | Body mass** | 5.723 ± 1.679 | 3.408 | **< 0.001** |
|  |  | Sex | 0.602 ± 0.406 | 1.484 | 0.138 |
|  |  | Body mass*Sex | -3.574 ± 2.153 | -1.660 | 0.100 |
|  |  | Chick ID: brood ID | 0.126 ± 0.357 | NA | NA |
|  | Frequency modulation<br>n = 2365 calls | Body mass** | 1.664 ± 0.703 | 2.368 | 0.018 |
|  |  | Sex | -0.100 ± 0.108 | -0.930 | 0.352 |
|  |  | Chick ID: brood ID | 0.057 ± 0.239 | NA | NA |
| Masked Lapwing | Call duration<br>n = 6683 calls | Body mass** | -0.299 ± 0.475 | -0.631 | 0.528 |
|  |  | Sex | 0.014 ± 0.013 | 1.024 | 0.306 |
|  |  | Chick ID: brood ID | 0.005 ± 0.071 | NA | NA |
|  | ICI*<br>n = 76 chicks | Body mass** | 1.980 ± 2.679 | 0.739 | 0.460 |
|  |  | Sex | 0.011 ± 0.084 | 0.134 | 0.893 |
|  |  | Chick ID: brood ID | 0.014 ± 0.117 | NA | NA |
|  | Entropy<br>n = 3096 calls | Body mass** | 2.283 ± 0.476 | 4.795 | **< 0.001** |
|  |  | Sex | -0.018 ± 0.015 | -1.270 | 0.204 |
|  |  | Chick ID: brood ID | 0.004 ± 0.061 | NA | NA |
|  | Minimum dominant frequency<br>n = 6683 calls | Body mass** | -3.269 ± 1.688 | -1.940 | 0.053 |
|  |  | Sex | -0.010 ± 0.051 | -0.190 | 0.850 |
|  |  | Chick ID: brood ID | 0.061 ± 0.248 | NA | NA |
|  | Dominant frequency range*<br>n = 6683 calls | Body mass** | 0.548 ± 0.556 | 0.986 | 0.324 |
|  |  | Sex | -0.069 ± 0.030 | -2.361 | 0.018 |
|  |  | Body mass*Sex | 2.210 ± 0.726 | 3.043 | **0.002** |
|  |  | Chick ID: brood ID | 0.004 ± 0.065 | NA | NA |
|  | Frequency modulation*<br>n = 6619 calls | Body mass** | 6.145 ± 0.951 | 6.46 | **< 0.001** |
|  |  | Sex | -0.084 ± 0.030 | -2.810 | **0.005** |
|  |  | Chick ID: brood ID | 0.016 ± 0.126 | NA | NA |

**Figure 1.**
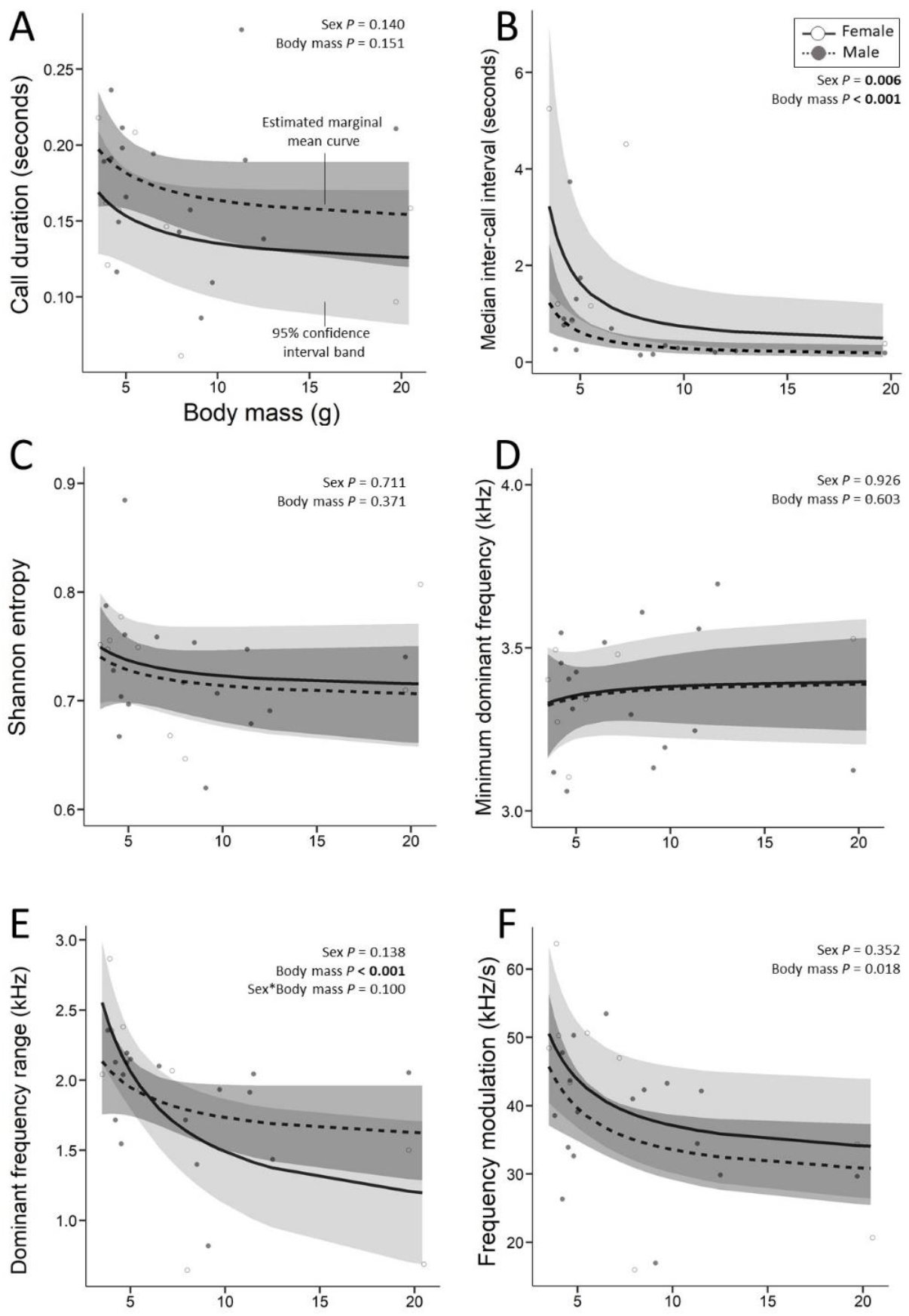
For plovers, call duration did not vary with body mass or sex (A), but inter-call intervals (ICIs) decreased with increasing body mass and were shorter in males (B). Entropy and minimum dominant frequency did not vary with body mass or sex (C−D). Dominant frequency range decreased with increasing body mass (E), but frequency modulation did not vary with body mass or sex (F). Estimated marginal means (hatched and solid lines) and 95% confidence intervals (grey shading) are shown.

For lapwings, frequency modulation was lower for males than for females (Table 1, Fig. 2). For both sexes, dominant frequency range decreased with increasing body mass; however, the decline was greater in males, resulting in a lower dominant frequency range than for females (Table 1; Fig. 2). As body mass increased, frequency modulation and entropy of lapwing calls decreased (Table 1; Fig. 2). Call duration, minimum dominant frequency and ICI were not predicted by sex or body mass.

**Figure 2.**
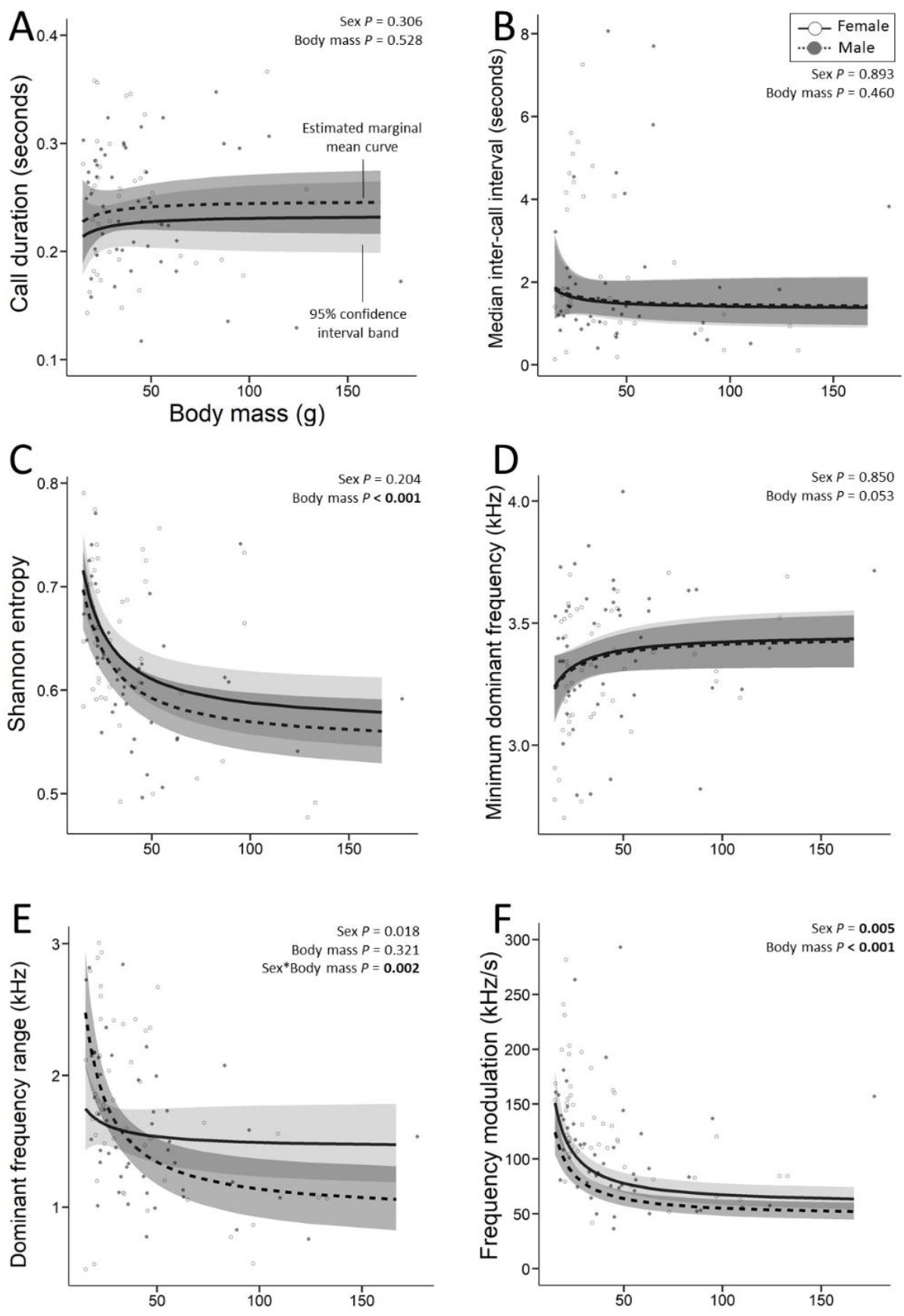
For lapwings, call duration, and inter-call intervals (ICIs) did not vary with body mass or sex (A-B), but entropy decreased with increasing body mass (C). Minimum dominant frequency did not vary with body mass or sex (D) but, for lapwings, dominant frequency range decreased with body mass and the decline was greater in males (E). Frequency modulation decreased with body mass and was lower in males (F). Estimated marginal means (hatched and solid lines) and 95% confidence intervals (grey shading) are shown.

## Discussion

Multiple call traits were associated with body mass; vocal traits changed gradually, as has been described previously in growing shorebirds (Adret 2012, Dragonetti et al. 2013), although repeated measurements of the same individuals are required to confirm this. For both study species, minimum dominant frequency was not associated with body mass, whereas dominant frequency range decreased with increasing body mass, consistent with maximum dominant frequency declining over growth in birds (Würdinger 1970, Adret 2012, Dragonetti et al. 2013). The decline in frequency modulation with increasing body mass in lapwings (and a similar but non-significant pattern for plovers) is consistent with the idea that frequency fluctuations within a call decrease with growth. Large bill size may impede the ability to quickly open and close the bill, therefore limiting the rate of frequency modulation (Podos 2001, Demery et al. 2021), though other explanations exist. Shorter ICIs were associated with heavier body masses in plovers, which might reflect the capacity of older, heavier, chicks to control airflow through vocal structures (Franz and Goller 2002, Zvonov 2011). We did not detect a change in call duration with age in either species (but see Adret 2012). Heavier lapwing chicks produced calls that were more tonal (i.e. reduced entropy), possibly associated with increased motor control of the upper vocal tract (Podos et al. 1995). As chicks age, variation in call structure may reflect attempts to utter adult-like calls, as described for other shorebirds (Adret 2012, Dragonetti et al. 2013).

For lapwings, frequency modulation was higher in females and, for plovers, male chicks had higher call rates than females. Compared to females, male Red-capped Plover chicks had shorter ICIs from hatching; cooing rates were also higher in male Pied Avocet chicks until the age of 9 months (Adret 2012). A faster repetition rate of distress calling might increase response intensity by adult defenders (Wheatcroft 2015), which could contribute to the higher survival of male over female chicks reported for several plover species (Sandercock et al. 2005, Pakanen et al. 2015, Saunders and Cuthbert 2015, Eberhart-Phillips et al. 2017; but see Lees et al. 2019). For lapwings, dominant frequency range declined with body mass more in males than in females, resulting in the development of a lower overall dominant frequency range in males. Frequency modulation was also lower in males than in females from hatching and throughout growth, suggesting differences in calls between the sexes at hatching. The sexes may differ in the size or rate of development in vocal anatomy, or the vocal control of these structures (Ballintijn and ten Cate 1997a, Gahr 2007, Volodin et al. 2015).

Overall, calls between the sexes were similar in most respects. Contextually, this study analysed distress calls (loud, harsh, and locatable; Sethi et al. 2012), which are likely under the influence of natural selection (Martin et al. 2011). Some non-distress vocalisations (e.g. contact calls) might communicate the caller’s sex, whereas distress calls may not (Austin et al. 2021). Even when a sex is “preferred” by parents, calls of the non-preferred sex are expected to evolve to be similar to those of the preferred sex (Austin et al. 2021). Furthermore, distress calls emitted by chicks may serve to communicate with siblings, other conspecifics, the predator, or some combination of these. Future studies should therefore investigate associations between shorebird chick calls and sex using the full repertoire of chick calls, and across species whose adult call repertoires and characteristics vary between sexes. We note our limited sample size, bias toward young chicks, and imbalance in sex ratio for plovers (34.6% were female) and suggest further study would be desirable. We also note that we did not repeatedly measure the same individuals, so cannot unambiguously exclude effects such as those associated with survival bias.

This research complies with the current laws of Australia and was conducted in accordance with protocols reviewed and approved by the Deakin University Animal Ethics Committee (Permit numbers B01 2018, B11 2017 and B12 2017) and the Department of Environment, Land, Water and Planning (Permits 10008437 and 10008619). The authors minimised impacts on chicks by only collecting data on days of suitable weather (e.g. no rain or extreme heat) and by generally avoiding recaptures. This project was supported by The Holsworth Wildlife Research Endowment and The Ecological Society of Australia. MAW was supported during write-up by the Beach Ecology and Conservation Hub (BEACH, Venus Bay). Thanks to Dr Peter Dann (Phillip Island Nature Parks) and Rangers at the Point Cook Coastal Park (Ron Cuthbert, Russell Brooks, Mark Cullen and Bernie McCarrick) for their assistance, to Ekaterina Ershova for her help in translating Russian material and to Rachel Adams (UTas) for bibliographic help.

## Author contributions

Methodology, K.N.K., M.A.W., D.R.W. and E.H.M.; Investigation, K.N.K.; Formal Analysis, K.N.K., D.R.W. and E.H.M.; Writing – Original Draft, K.N.K.; Writing – Review and Editing, K.N.K., M.A.W., D.R.W. and E.H.M.; Funding Acquisition, K.N.K.; Supervision, M.A.W.

## Data availability statement

The datasets generated during and/or analysed during the current study are available in the figshare repository via: 10.6084/m9.figshare.14813379

## SUPPORTING INFORMATION

Additional supporting information may be found online in the Supporting Information section at the end of the article.

### Supplemental Methods – Acoustic Analysis

Details of the quantification of acoustic variables

**Figure S1.**
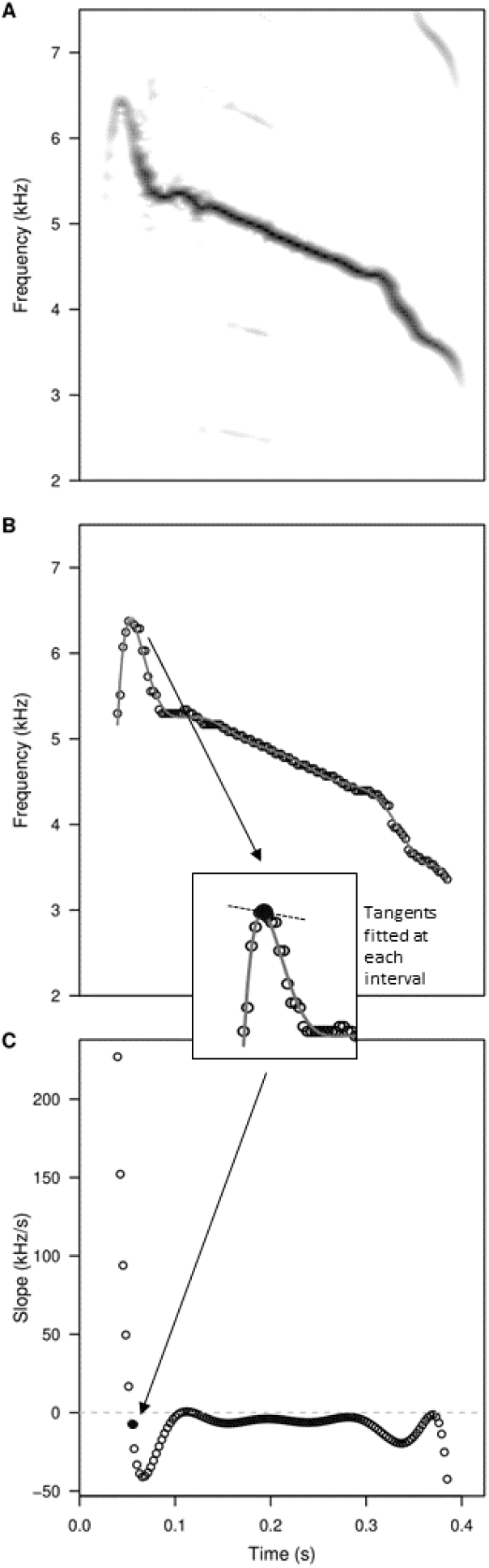
To quantify the structure of Masked lapwing calls we first inspected the spectrogram of each call (A). Calls were then divided into contiguous 2.9-ms time bins, and a mean power spectrum calculated for each bin. We measured the dominant frequency of each bin from the corresponding mean spectrum and plotted it as a function of time (B; open circles). We regressed dominant frequency values on time using a series of polynomial regressions (first to twelfth order), and superimposed the best-fitting regression on the dominant frequency values (solid line; B). We calculated the slope of the tangent to the selected polynomial for every 2.9-ms interval and plotted it as a function of time (open circles; C). The inset shows an example of a tangent (solid circle and hatched line), for the 2.9-ms interval marked in other panels (B and C) by a solid black circle. We used the range in slopes to characterise the magnitude of frequency modulation throughout the call. The grey hatched line indicates a slope of zero (C).

**Figure S2.**
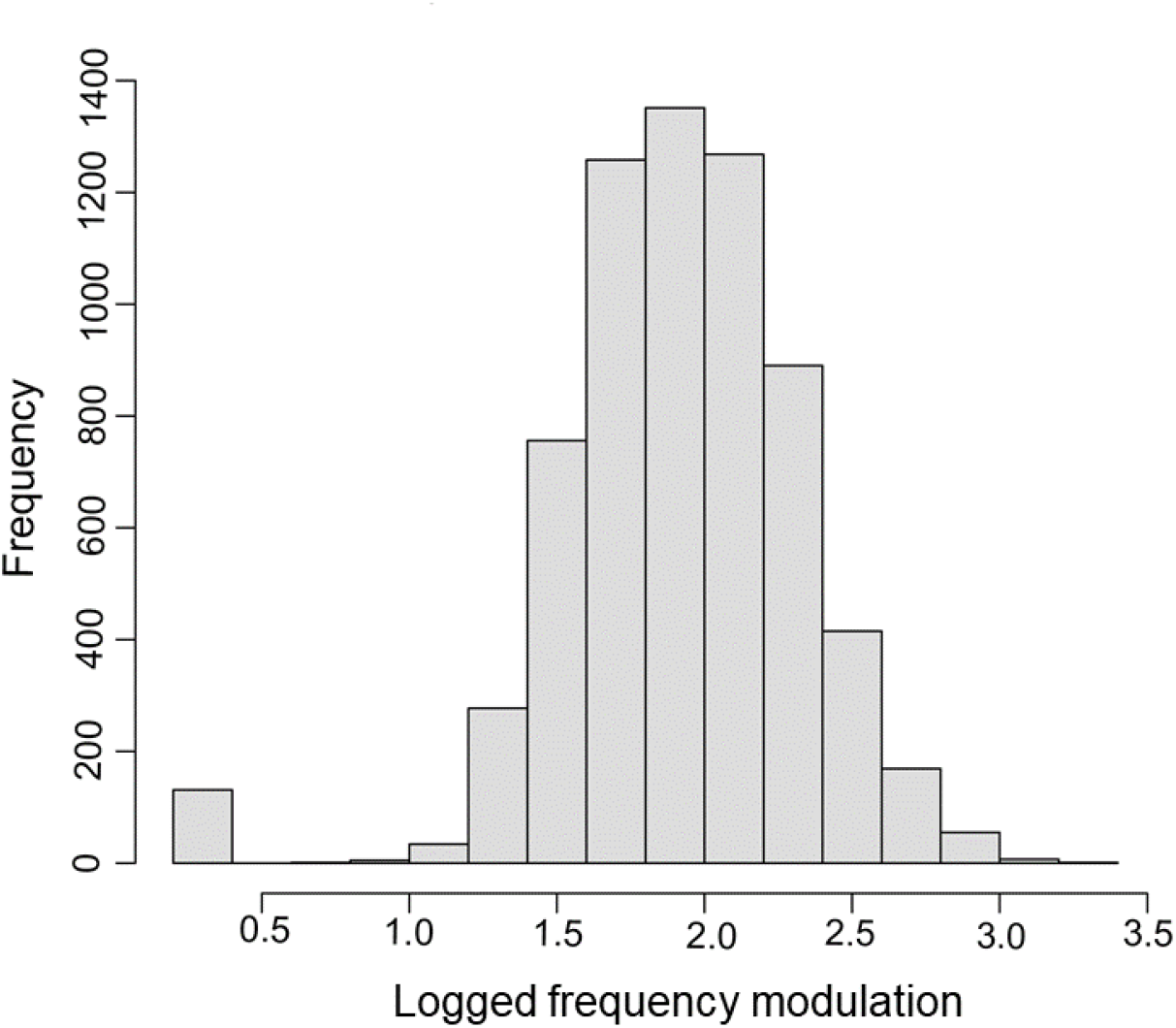
The frequency distribution of log-transformed frequency modulation (rate of change in dominant frequency) for lapwing chick calls.

**Figure S3.**
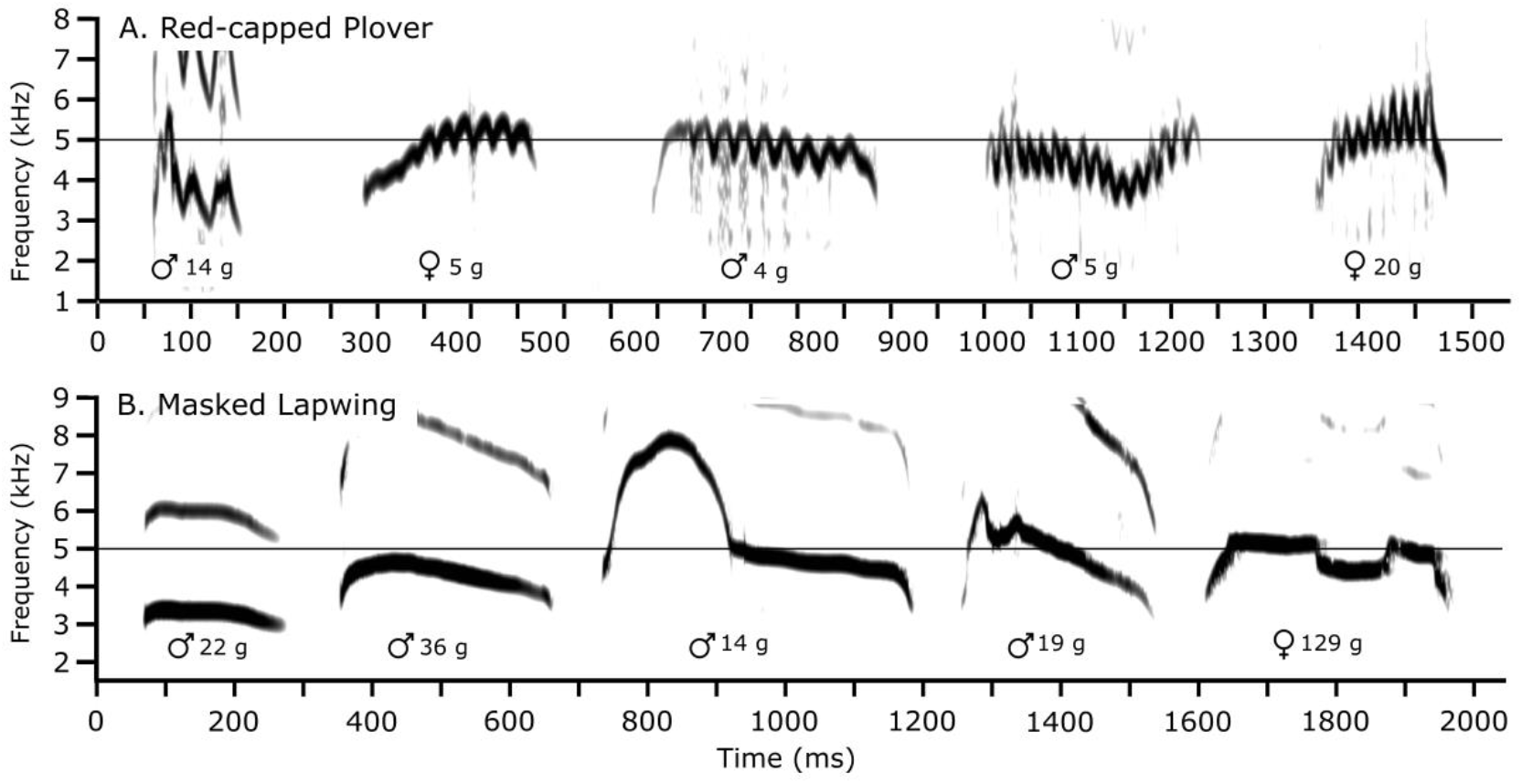
Chick distress calls differed substantially between Red-capped Plover (A) and Masked Lapwing (B), particularly in the presence and extent of frequency modulation (FM). Calls are ordered left-to-right by rate of FM for plovers and by increasing overall complexity for lapwings.

**Table S1.** Body measurements for all Red-capped Plover chicks within this study.

| Date | Known age | Recording number | Chick ID | Sex | Brood number | Body mass | Head bill | Bill | Tarsus | Tarsus and toe | Recapture |
| --- | --- | --- | --- | --- | --- | --- | --- | --- | --- | --- | --- |
| 3/01/2019 | 1.0 | R350 | chick 2 | male | 1 | 4.5 | 22.6 | 17.6 | 19.42 | 33 |  |
| 20/01/2019 |  | R362 | chick 2a | female | 3 | 4 | 24.12 | 7.6 | 19.02 | 34.5 |  |
| 1/02/2019 |  | R379 | 3691822 | male | 4 | 11.3 | 29.43 | 10.78 | 22.35 | 40 |  |
| 1/02/2019 |  | R380 + 381 | 3691823 | male | 5 | 8.5 | 29.41 | 10.62 | 23.96 | 41 |  |
| 4/03/2019 |  | Z240 | 3691820 | male | 18 | 6.3 | 25.9 | 8.09 | 19.66 | 38 |  |
| 1/02/2019 | 1.0 | R382 | 3691824 | female | 6 | 5.5 | 24.2 | 7.59 | 16.35 | 35 |  |
| 22/03/2019 |  | R410 | 3691820 | male | 18 | 14.5 | 32.64 | 9.83 | 23.55 | 42 | Y |
| 1/02/2019 | 1.0 | R383 | 3691825 | female | 6 | 3.5 | 24.89 | 8.16 | 18.62 | 35 |  |
| 11/02/2019 | 1.0 | R391 | 3691827 | male | 8 | 5 | 20.71 | 6.25 | 16.5 | 31 |  |
| 11/02/2019 |  | R394 | 3691835 | male | 11 | 4.2 | 23.89 | 7.82 | 17.8 | 34 |  |
| 11/02/2019 |  | R393 | 3691829 | male | 9 | 19.7 | 35.24 | 12 | 24.98 | 45 |  |
| 18/02/2019 |  | Z232 | 3691831 | female | 14 | 8 | 27.59 | 9.3 | 20 | 40 |  |
| 11/02/2019 |  | R392 | 3691828 | male | 12 | 5.1 | 23.37 | 7.31 | 18.65 | 36 |  |
| 18/02/2019 |  | Z233+Z234 | 3691832 | female | 14 | 20.5 | 35.7 | 12.88 | 26.02 | 44.5 |  |
| 4/03/2019 |  | Z238 | 3691828 | male | 12 | 21.8 | 33.73 | 10.91 | 24.99 | 45 | Y |
| 18/02/2019 |  | Z230 +Z231 | 3691833 | female | 13 | 19.7 | 35.13 | 11.31 | 25.42 | 42.5 |  |
| 18/02/2019 |  | Z229 | 3691834 | female | 12 | 7.2 | 26.52 | 8.5 | 20.75 | 38.5 | Y |
| 18/02/2019 |  | Z235 | 3691836 | male | 15 | 11.5 | 29.96 | 10.1 | 20.44 | 39.5 |  |
| 11/02/2019 |  | R396 | 3691834 | female | 12 | 5.3 | 23.68 | 7.3 | 19.3 | 46 |  |
| 25/02/2019 |  | Z236 | 3691828 | male | 12 | 12.5 | 30.51 | 9.61 | 21.79 | 41 | Y |
| 4/03/2019 |  | Z239 | 3693837 | male | 17 | 7.9 | 26.88 | 8.79 | 22.6 | 42 |  |
| 8/03/2019 | 1.0 | R405 | 3691839 | male | 19 | 3.8 | 23.83 | 7.74 | 19.15 | 46 |  |
| 8/03/2019 | 1.0 | R404 | 3691838 | male | 19 | 3.6 | 25.24 | 7.55 | 18.88 | 36 |  |
| 8/03/2019 |  | R406 | 3691840 | male | 18 | 9.7 | 27.34 | 8.85 | 19.98 | 38 | Y |
| 14/03/2019 |  | R407 | 3691820 | male | 18 | 9.1 | 29.14 | 10.8 | 20.46 | 39.5 | Y |
| 14/03/2019 |  | R408 | 3691840 | male | 18 | 8.2 | 29.23 | 9.19 | 21.67 | 44 | Y |
| 14/03/2019 | 7.0 | R409 | 3691838 | male | 19 | 4.8 | 26.7 | 8.15 | 19.15 | 37 | Y |
| 22/03/2019 |  | R411 | 3691840 | male | 18 | 14.8 | 31.85 | 11.07 | 22.04 | 41 | Y |
| 16/12/2019 | 1.0 | Z255 | 03691814 | male | 24 | 4.6 | 21.79 | 7.1 | 16.51 | 33 |  |
| 16/12/2019 | 1.0 | Z254 | 03691815 | female | 23 | 4.6 | 22.05 | 7.02 | 17.34 | 33 |  |
| 16/12/2019 | 1.0 | Z252 | 03691817 | male | 22 | 6.5 | 24.22 | 7.65 | 17.92 | 34.5 |  |
| 16/12/2019 | 1.0 | Z251 | 03691818 | female | 22 | 3.9 | 23.35 | 7.78 | 17.42 | 35 |  |
| 16/12/2019 | 1.0 | Z247 | 03691845 | male | 20 | 4.8 | 22.98 | 17.94 | 16.83 | 34 |  |
| 16/12/2019 | 1.0 | Z250 | 03691846 | male | 21 | 4.2 | 23.3 | 7.52 | 18.3 | 36 |  |
| 16/12/2019 |  | Z247 | 03691845 | male | 20 | 4.8 | 22.98 | 17.94 | 16.83 | 34 |  |
| 16/12/2019 |  | Z250 | 03691846 | male | 21 | 4.2 | 23.3 | 7.52 | 18.3 | 36 |  |
| 16/12/2019 |  | Z251 | 03691818 | female | 22 | 3.9 | 23.35 | 7.78 | 17.42 | 35 |  |
| 16/12/2019 |  | Z252 | 03691817 | male | 22 | 6.5 | 24.22 | 7.65 | 17.92 | 34.5 |  |
| 16/12/2019 |  | Z254 | 03691815 | female | 23 | 4.6 | 22.05 | 7.02 | 17.34 | 33 |  |
| 16/12/2019 |  | Z255 | 03691814 | male | 24 | 4.6 | 21.79 | 7.1 | 16.51 | 33 |  |

**Table S2.** Body measurements for all Masked Lapwing chicks within this study.

| Date | Known age | Recording number | Chick ID | Sex | Brood number | Body mass | Head bill | Bill | Tarsus | Tarsus and toe |
| --- | --- | --- | --- | --- | --- | --- | --- | --- | --- | --- |
| 2/08/2018 | 1.0 | R63 + R64 + Z71 | 083- 24303 | male | 1 | 18.2 | 37.7 | 11.89 | 27.51 | 59 |
| 2/08/2018 | 1.0 | R65 | 083- 24304 | male | 1 | 22.4 | 36 | 12.32 | 32.19 | 57.1 |
| 2/08/2018 | 1.0 | R66 | 083- 24305 | female | 1 | 22.5 | 35.41 | 12.01 | 31.09 | 58 |
| 2/08/2018 | 1.0 | R67 | 083- 24306 | male | 2 | 22.4 | 35.09 | 11.02 | 28.76 | 55.5 |
| 2/08/2018 | 1.0 | R68 | 083- 24307 | female | 2 | 21.1 | 36.07 | 11.31 | 30.33 | 59 |
| 2/08/2018 | 1.0 | R69 | 083- 24308 | female | 2 | 22.2 | 35.6 | 11.65 | 28.99 | 58 |
| 2/08/2018 | 1.0 | R71 | 083- 24310 | male | 3 | 17 | 32.17 | 9.8 | 27.82 | 52 |
| 3/08/2018 | 1.0 | R72 | 083- 24311 | male | 4 | 21.5 | 33.2 | 10.47 | 28.3 | 55.5 |
| 3/08/2018 |  | R76 | 083- 24312 | female | 5 | 45.4 | 47.29 | 19.21 | 33.49 | 64 |
| 3/08/2018 | 2.0 | R84 | 083- 24313 | female | 6 | 21.8 | 37.98 | 13.79 | 21.15 | 59.5 |
| 3/08/2018 | 2.0 | R78 + R81 | 083- 24314 | male | 6 | 19.4 | 37.92 | 14.01 | 30.26 | 60 |
| 3/08/2018 | 2.0 | R83 | 083- 24315 | female | 6 | 20.7 | 37.06 | 13.8 | 29.84 | 58 |
| 3/08/2018 | 2.0 | R79 + R80 + R82 | 083- 24316 | male | 6 | 19.5 | 36.81 | 12.95 | 29.64 | 59 |
| 3/08/2018 |  | R85 | 083- 24317 | female | 7 | 23.9 | 42.95 | 16.42 | 29.66 | 60 |
| 3/08/2018 |  | R86 | 083- 24318 | female | 7 | 23.1 | 41.6 | 15 | 30 | 59 |
| 3/08/2018 |  | R87 | 083- 24319 | male | 9 | 28 | 44.43 | 17.45 | 34.02 | 65 |
| 3/08/2018 |  | R88 | 083- 24320 | female | 9 | 29 | 44.06 | 18.66 | 32.88 | 64 |
| 3/08/2018 |  | R89 | 083- 24321 | female | 10 | 22.5 | 35.52 | 12.44 | 28.89 | 58 |
| 3/08/2018 | 2.0 | R90 | 083- 24322 | female | 11 | 22.5 | 34.49 | 10.58 | 28.6 | 58 |
| 3/08/2018 | 2.0 | R91 | 083- 24323 | male | 11 | 21.5 | 36.6 | 11.91 | 31.11 | 60.5 |
| 3/08/2018 |  | R93 + R94 | 083- 24325 | male | 12 | 95 | 55.09 | 22.79 | 44.94 | 80 |
| 3/08/2018 | 7 or 8 | R 96 + R100 | 083- 24326 | male | 13 | 22.3 | 41.09 | 15.3 | 31.74 | 63 |
| 3/08/2018 | 7 or 8 | R 97 +R99 | 083- 24327 | male | 13 | 25.2 | 40.75 | 13.99 | 32.08 | 61 |
| 3/08/2018 | 7 or 8 | R98 | 083- 24328 | male | 13 | 17.9 | 41.69 | 15.6 | 32.14 | 60.5 |
| 3/08/2018 | 7 or 8 | R101 | 083- 24329 | female | 13 | 26.5 | 42.09 | 14.9 | 32.21 | 60 |
| 3/08/2018 | 10 or 11 | R102 | 083- 24330 | female | 14 | 37.4 | 46.45 | 17.85 | 36.6 | 68 |
| 3/08/2018 | 10 or 11 | R103 + R106 | 083- 24331 | male | 14 | 33.4 | 42.91 | 13.83 | 32.84 | 62 |
| 3/08/2018 | 10 or 11 | R104 | 083- 24332 | female | 14 | 46.8 | 47.42 | 17.15 | 37.28 | 69.5 |
| 3/08/2018 | 10 or 11 | R105 | 083- 24333 | male | 14 | 44.9 | 47.3 | 17.48 | N/A | 68 |
| 6/08/2020 | 5 or 6 | R107 | chick 1d | male | 16 | 15.5 | 36.35 | 12.51 | 28.19 | 53 |
| 9/08/2018 |  | R118 | 083- 24338 | male | 17 | 63 | 51 | 21.58 | 39.18 | 72 |
| 9/08/2018 |  | R119 + R121 | 083- 24339 | female | 17 | 73 | 52.75 | 22.67 | 43.7 | 76 |
| 9/08/2018 |  | R120 | 083- 24340 | female | 17 | 65 | 52.91 | 23.76 | 40.65 | 73 |
| 9/08/2018 |  | R123 | 083- 24341 | female | 18 | 18.6 | 37.19 | 12.85 | 29.97 | 59 |
| 9/08/2018 |  | R126 | 083- 24342 | female | 18 | 20.7 | 37.43 | 12.39 | 31.69 | 58 |
| 9/08/2018 |  | R132 + R133 | 083- 24345 | female | 19 | 24.3 | 37.9 | 13.8 | 31.79 | 62 |
| 9/08/2018 |  | R134 | 083- 24346 | male | 19 | 21.1 | 37.33 | 12.53 | 31.19 | 58 |
| 9/08/2018 |  | R140 | 083- 24348 | female | 20 | 44.6 | 48.4 | 17 | 37.52 | 69 |
| 9/08/2018 |  | R143 | 083- 24350 | female | 20 | 39.55 | 46.72 | 17.33 | 34.26 | 68 |
| 9/08/2018 | 3.0 | R145 | 083-36151 | female | 21 | 23 | 39.11 | 14.67 | 34 | 62 |
| 9/08/2018 | 3.0 | R146 | 083-36152 | female | 21 | 21.2 | 37.46 | 13.14 | 31.47 | 59.5 |
| 9/08/2018 |  | R147 | 083-36153 | male | 22 | 37 | 46.26 | 17.93 | 34.59 | 64 |
| 9/08/2018 |  | R148 | 083-36154 | male | 22 | 36.2 | 45 | 18.89 | 34.29 | 63 |
| 9/08/2018 |  | R149 | 083-36155 | female | 22 | 34 | 44.79 | 18.35 | 35.83 | 63 |
| 10/08/2018 | 1 to 3 | R154 | 083-36158 | female | 24 | 21.2 | 35.81 | 11.39 | 31 | 59 |
| 10/08/2018 | 1 to 3 | R157 R158 | 083-36159 | male | 24 | 19.5 | 37.45 | 12.4 | 29.66 | 56.5 |
| 10/08/2018 | 1 to 3 | R159 | 083-36160 | female | 24 | 20.1 | 34.65 | 10.68 | 30.22 | 56 |
| 10/08/2018 |  | R160 | 083-36161 | female | 25 | 19.4 | 38.43 | 14.25 | 32.69 | 61.5 |
| 10/08/2018 |  | R161 | 083-36162 | male | 25 | 18.5 | 38.31 | 14.71 | 30.04 | 58 |
| 10/08/2018 |  | R162 | 083-36163 | female | 25 | 15.2 | 39.83 | 15.28 | 34.69 | 63.5 |
| 16/08/2018 |  | R194 + 195 | chick 1 | male | 27 | 55 | 50.62 | 22.55 | 42.15 | 75 |
| 16/08/2018 |  | R194 | chick 2 | female | 27 | 53.9 | 48.25 | 19.5 | 39.3 | 70 |
| 23/08/2018 |  | R208 + 209 | 083-36166 | male | 29 | 43.5 | 45.68 | 16.62 | 35.89 | 65 |
| 23/08/2018 |  | R210 | 083-36167 | male | 29 | 49 | 46.96 | 19.24 | 36 | 63.5 |
| 23/08/2018 |  | R211 | 083-36168 | female | 30 | 97 | 54.49 | 21.15 | 47.35 | 81 |
| 23/08/2018 |  | R212 | 083-36169 | female | 30 | 97 | 53.81 | 21.3 | 46.83 | 80 |
| 23/08/2018 |  | R213 | 083-36170 | male | 31 | 36 | 43.25 | 15.61 | 33.59 | 62 |
| 23/08/2018 |  | R214 | 083-36171 | male | 32 | 87 | 54.79 | 24.55 | 44.7 | 80 |
| 23/08/2018 |  | R215 | 083-36172 | female | 33 | 30.3 | 42.46 | 15 | 34.45 | 61 |
| 23/08/2018 |  | R216 | 083-36174 | male | 33 | 26.5 | 41.19 | 15 | 33.6 | 62 |
| 23/08/2018 |  | R217 | 083-36175 | female | 33 | 33.5 | 42.46 | 15.78 | 35.01 | 64 |
| 23/08/2018 | 3.0 | R221 | 083-36177 | female | 34 | 18.5 | 37.69 | 12.2 | 29.48 | 58 |
| 23/08/2018 | 3.0 | R222 | 083-36178 | female | 34 | 20.2 | 38.2 | 13.05 | 30 | 57 |
| 23/08/2018 | 10.0 | R223 | 083-36179 | male | 35 | 25.5 | 40.26 | 15.29 | 31.45 | 57 |
| 23/08/2018 | 10.0 | R224 | 083-36180 | male | 35 | 35.3 | 44.81 | 17.4 | 33.7 | 64.5 |
| 24/08/2018 |  | R228 | 083-36181 | male | 36 | 58.8 | 48.97 | 19.39 | 38.07 | 71 |
| 24/08/2018 |  | R229 | 083-36182 | female | 36 | 109 | 59.14 | 26.58 | 48.16 | 85 |
| 24/08/2018 |  | R230 | 083-36183 | male | 36 | 110 | 58.08 | 25.71 | 49.65 | 83.5 |
| 24/08/2018 | 14.0 | R231 | 083-36184 | male | 37 | 56.1 | 47.83 | 17.49 | 37.34 | 68.5 |
| 24/08/2018 |  | R232 | 083-36185 | male | 38 | 83 | 53.84 | 22.21 | 35 | 80 |
| 24/08/2018 |  | R233 | 083-36186 | female | 38 | 47 | 48.78 | 20.5 | 34.85 | 65 |
| 12/09/2018 | 13 to 14 | R318 | chick 1a | female | 39 | 41.8 | 46.3 | 17.92 | 35.11 | 67 |
| 12/09/2018 |  | R321 | chick 1b | female | 41 | 29 | 41.3 | 15.75 | 32.68 | 59 |
| 12/09/2018 |  | R322 | chick 1c | male | 42 | 30 | 43.21 | 18.42 | 34.49 | 62 |
| 12/09/2018 |  | R323 | chick 2a | male | 42 | 24.5 | 42.37 | 15.76 | 33.8 | 61.5 |
| 20/09/2018 |  | T56 | 083-36188 | female | 44 | 35 | 42.81 | 16.81 | 32.84 | 61 |
| 20/09/2018 |  | T57 | 083-36189 | female | 44 | 28.7 | 42.02 | 17.3 | 22.33 | 61.5 |
| 20/09/2018 |  | T58 | 083-36190 | male | 45 | 32.5 | 45 | 18.35 | 32.19 | 61 |
| 20/09/2018 |  | T59 + 60 | 083-36173 | female | 46 | 44 | 47.41 | 18.72 | 36.13 | 66.5 |
| 20/09/2018 |  | T61, 62, 63 + 64 | 083-36191 | male | 46 | 62.8 | 49.7 | 19.38 | 40.22 | 71 |
| 20/09/2018 |  | T65 | 083-36192 | male | 46 | 41 | 46.18 | 17.04 | 34.79 | 66 |
| 20/09/2018 |  | T69 | 083-36196 | female | 48 | 50.5 | 46.54 | 17.91 | 36.22 | 64.5 |
| 20/09/2018 |  | T67 + 70 | 083-36194 | male | 48 | 48.3 | 47.38 | 19.44 | 37.72 | 70 |
| 20/09/2018 |  | T68 | 083-36195 | male | 48 | 47.6 | 47.63 | 17.76 | 37.49 | 67 |
| 20/09/2018 |  | T71 | 083-36197 | male | 49 | 177 | 62.84 | 28.29 | 59.84 | 97 |
| 20/09/2018 |  | T75 | 083-27551 | female | 50 | 86 | 55.43 | 24.45 | 47.84 | 81.5 |
| 20/09/2018 |  | T72 | 083-36198 | male | 50 | 55.4 | 49.3 | 20.71 | 39.3 | 69 |
| 20/09/2018 |  | T73 | 083-36199 | male | 50 | 89 | 52.61 | 21.25 | 46.68 | 81 |
| 21/09/2018 |  | T76 | 083-27552 | female | 51 | 129 | 59.96 | 26.75 | 52.12 | 92 |
| 21/09/2018 |  | T77 | 083-27553 | male | 51 | 124 | 59.5 | 27.34 | 55.69 | 94 |
| 21/09/2018 |  | T78 | 083-27554 | male | 52 | 39.7 | 45 | 16.92 | 36.64 | 68 |
| 21/09/2018 |  | T79 | 083-27555 | male | 53 | 45.2 | 44.58 | 15.5 | 34.99 | 65.5 |
| 21/09/2018 |  | T81 | 083-27557 | male | 55 | 31.4 | 45.38 | 18.49 | 33.32 | 61 |
| 21/09/2018 |  | T82 | 083-27558 | male | 56 | 44.9 | 47.5 | 18.54 | 36.98 | 67 |
| 21/09/2018 |  | T83 | 083-27559 | female | 57 | 133 | 58.4 | 25.23 | 51.11 | 88 |
| 21/09/2018 |  | T84 | 083-27560 | male | 58 | 49.7 | 45.41 | 18.25 | 35 | 66 |

**Table S3.** Results of the generalised linear mixed model investigating the relationship between frequency modulation of lapwing calls with outliers removed (> 1.5 times the interquartile ranges below the first or above the third quartile) and body mass (reciprocal transformation) and sex (reference category = female). We specified a Toeplitz covariance structure, and chick identity nested within brood identity was included as the random effect for all analyses. Estimates are presented as estimates of coefficients ± standard error for fixed effects, and variance ± standard deviation for the random effect of Identification nested within Brood ID. The probability distribution was Gaussian and we used an identity link. The response variable was logarithmically transformed. * = to aid interpretation, note that the reciprocal transformation reflects the sign of coefficients.

| Species | Response variable | Model terms | Estimates | *z* | *P* |
| --- | --- | --- | --- | --- | --- |
| Masked Lapwing | Frequency modulation<br>*n* = 6456 calls | Body mass* | 5.626 ± 0.873 | 6.450 | **< 0.001** |
| | | Sex | -0.091 ± 0.027 | -3.330 | **< 0.001** |
| | | Identification: Brood ID | 0.014 ± 0.119 | NA | NA |

**Table S4.** Results of the generalised linear mixed model investigating the relationships of body mass (reciprocal transformation), sex (reference category = female), and call saturation (reference category = not saturated) with call entropy for both species. We specified a Toeplitz covariance structure, and chick identity nested within brood identity was included as the random effect for all analyses. Estimates are presented as estimates of coefficients ± standard error for fixed effects, and variance ± standard deviation for the random effect of identification nested within brood ID. The probability distribution was Gaussian and we used an identity link. * = logarithmically transformed data. ** = to aid interpretation, note that the reciprocal transformation reflects the sign of coefficients.

| Species | Response variable | Model terms | Estimates | z | P |
| --- | --- | --- | --- | --- | --- |
| Red-capped Plover | Entropy*<br>n = 2365 calls | Body mass** | -0.069 ± 0.047 | -1.455 | 0.146 |
| | | Sex | 0.003 ± 0.007 | 0.413 | 0.679 |
| | | Saturation | -0.013 ± 0.001 | -16.810 | **< 0.001** |
| | | Identification: Brood ID | 0.000 ± 0.017 | NA | NA |
| Masked Lapwing | Entropy<br>n = 6456 calls | Body mass** | 1.810 ± 0.373 | 4.850 | **< 0.001** |
| | | Sex | -0.003 ± 0.011 | -0.280 | 0.780 |
| | | Saturation | -0.020 ± 0.002 | -9.610 | **< 0.001** |
| | | Identification: Brood ID | 0.003 ± 0.055 | NA | NA |

## SUPPORTING INFORMATION

### Supplemental Methods – Acoustic Analysis

Using Raven Pro 1.6.1 (www.birds.cornell.edu/raven), we viewed each recording as a waveform and spectrogram (plover, fast Fourier transform (FFT) size = 512 samples; lapwing, FFT size = 1024 samples; hamming window and 87.5% overlap for both species). Frequency modulation was rapid in many plover calls, so we specified a shorter FFT window for that species to resolve frequency changes better; this was not necessary for lapwing calls, which featured little frequency modulation (Fig. S3; definition below). Using spectrograms as guides, we manually marked the approximate beginning and end of every call of each chick (or its identifiable sibling) on the waveform. Using these time records, we then opened each call in R (R Core Team 2021), plus 30 ms before and after the call, using the seewave and tuneR packages (Sueur et al. 2008, Ligges et al. 2018). We applied a 1000 Hz high-pass filter, which reduced low-frequency background noise without affecting call structure, and normalised calls to 0 dB. For lapwing calls, we adjusted the start and end times using a more objective approach based on an amplitude threshold, which defined the start of calls as the point where amplitude exceeded 15% of the maximum amplitude for 2 ms and the end of calls as the point where amplitude fell below 15% for 2 ms. We plotted the new start and end times on waveforms and spectrograms (settings as above) to ensure that values derived from the amplitude threshold approach were not based on artifacts. In a few cases (40 of 6835 calls), high background noise masked the start and end times, so we relied on our original (manually determined) times for those calls. For plover calls, low signal-to-noise ratios precluded the use of the amplitude threshold approach in many calls, so we used the manually selected start and end times.

We divided each call into a series of contiguous 2.9-ms time bins, and, for each bin, measured the Shannon spectral entropy and dominant frequency from a mean power spectrum (settings as per the corresponding spectrogram listed above). Entropy is a measure of tonal purity that approaches 0 for pure tones and 1 for white noise, and dominant frequency is the frequency of maximum amplitude, excluding harmonics and background noises. We validated our measures of dominant frequency by plotting them against time and comparing the resulting plot to the corresponding spectrogram (Fig. S1). In some cases, measures of dominant frequency, particularly at the start or end of the call, were based on background noise rather than the call itself. If these measurement errors were limited to one or two time bins (i.e. 2.9 or 5.8 ms), we excluded those bins from subsequent analyses; if they occurred in more than two time bins, or in the middle of the call, we excluded the entire call from subsequent analyses (we thereby excluded 235 of 2600 plover calls, and 153 of 6835 lapwing calls).

